# Inhibition of hypoxia-inducible factors suppresses subretinal fibrosis

**DOI:** 10.1101/2023.12.12.571193

**Authors:** Chiho Shoda, Deokho Lee, Yukihiro Miwa, Satoru Yamagami, Hiroyuki Nakashizuka, Kazumi Nimura, Kazutoshi Okamoto, Hirokazu Kawagishi, Kazuno Negishi, Toshihide Kurihara

## Abstract

Age-related macular degeneration (AMD) is a common cause of vision loss. The aggressive form of AMD is associated with ocular neovascularization and subretinal fibrosis, representing a responsive outcome against neovascularization mediated by epithelial-mesenchymal transition of retinal pigment epithelium cells. A failure of the current treatment (anti-vascular endothelial growth factor therapy) has also been attributed to the progression of subretinal fibrosis. Hypoxia-inducible factors (HIFs) increase gene expressions to promote fibrosis and neovascularization. HIFs act as a central pathway in the pathogenesis of AMD. HIF inhibitors may suppress ocular neovascularization. Nonetheless, further investigation is required to unravel the aspects of subretinal fibrosis. In this study, we used RPE-specific HIFs or von Hippel-Lindau (VHL, a regulator of HIFs) conditional knockout (cKO) mice, along with pharmacological HIF inhibitors, to demonstrate the suppression of subretinal fibrosis. Fibrosis was suppressed by treatments of HIF inhibitors, and similar suppressive effects were detected in RPE-specific *Hif1a*/*Hif2a-* and *Hif1a*-cKO mice. Promotive effects were observed in RPE-specific *Vhl*-cKO mice, where fibrosis-mediated pathologic processes were evident. Marine products’ extracts and their component taurine suppressed fibrosis as HIF inhibitors. Our study shows critical roles of HIFs in the progression of fibrosis, linking them to the potential development of therapeutics for AMD.

## Introduction

Age-related macular degeneration (AMD) is a major cause of vision loss among older people. AMD includes two dry and wet forms (1), depending on the existence of geographic atrophy and choroidal neovascularization (CNV) (2, 3). Wet AMD is more aggressive than dry AMD, causing rapid and severe vision loss. AMD can be characterized by the associated symptoms: early dry AMD (no specific symptoms), intermediate dry AMD (no symptoms or mild blurriness in the central vision), late AMD (wet or dry; noticing straight wavy or crooked lines, a blurry area with blank spots, or trouble seeing in low lighting).

Macular neovascularization (MNV/CNV) originates from the choroid through a break in the Bruch’s membrane into the sub-retinal pigment epithelium (RPE) or subretinal space, which increases the possibility of hemorrhage and exudative alterations and progresses to subretinal fibrosis. Subretinal fibrosis is characterized by intricate interactions within subretinal lesions between cellular components and inflammatory mediators, including transforming growth factor (TGF)-β and platelet-derived growth factor (PDGF) (4, 5). The extracellular matrix (ECM) can be reconstructed by fibrosis to form a subretinal scar. The prominent components of the ECM in subretinal fibrosis are proteins in the collagen family (especially, types I or IV) and fibronectin with minor amounts of collagen types III, V, and VI (6). This pathophysiologic process can lead to severe visual loss. The epithelial-mesenchymal transition (EMT) of RPE cells has been known a critical pathologic contributor during the development of subretinal fibrosis (7). RPE cells form a functional unit with photoreceptors in the outer retina and the choroid for visual function, and dysfunctions of the RPE have been considered as an initiating or early factor for the development of AMD (8). Thus, targeting RPE cells has been suggested as a promising therapeutic strategy in AMD.

Hypoxia-inducible factors (HIFs) are dimeric protein complexes that have important roles in response to low oxygen concentrations in tissues. HIFs are constantly synthesized at the physiological level (9). However, HIFs are rapidly hydroxylated by prolyl hydroxylase and are then bound to von Hippel-Lindau tumor-suppressor protein (VHL) for proteasome-mediated degradation under normoxic conditions (10). However, under hypoxic conditions, accumulated HIFs (undegraded) induce the expression of various genes, including pyruvate dehydrogenase kinase (*PDK1*), bcl-2 interacting protein 3 (*BNIP3*), vascular endothelial growth factor (*VEGF*), and glucose transporter protein type 1 (*GLUT1*), as strong master regulators of oxygen homeostasis depending on the cell type (11). Although those responses may solve the transient hypoxic condition, accumulating evidence has demonstrated that HIFs could be chronically involved in the pathogenesis of ocular neovascularization (the retina and the choroid) (12–14). Furthermore, HIFs have been gradually considered to be involved in subretinal fibrosis and EMT processes in RPE cells (15–17). Hence, pharmacologically or genetically regulating HIFs may enable novel treatment of wet AMD with subretinal fibrosis as well as MNV.

In this study, we aimed to understand the pathological role of HIFs in subretinal fibrosis development in various genetically engineered experimental mice. We investigated whether novel HIF inhibitors (extracts from natural marine products (18) and the effector molecule taurine from those products identified in our HIF inhibitor screening process) could have therapeutic effects on suppressing subretinal fibrosis in these experimental models. Accordingly, we suggest a novel therapeutic approach for wet AMD.

## Results

### Modulation of HIFs in the RPE affects fibrosis formation in adult mice

Although accumulating evidence has shown that HIFs may be the critical contributors for the development of MNV or further fibrosis (19–22), it remains unclear whether HIF-1α, HIF-2α, or both are the main contributor(s). Therefore, we produced RPE-specific *Hifs*-conditional knockout (cKO) mice (23) (*Hif1a*^f/f^/*Hif2a*^f/f^, *Hif1a*^f/f^, or *Hif2a*^f/f^; Best1-Cre^tg/-^) to examine which HIF gene(s) are mainly involved (**Fig. 1**). Formation of fibrosis has been reported at the chronic stage of laser-induced CNV in experimental models (24–27). Using collagen I α1 immunostaining, we found that fibrosis formed where CNV (stained by IB4) chronically remained. The fibrosis volume was considerably reduced by double cKO of *Hif* genes (**Fig. 1A**), and although *Hif2a*-cKO only produced a decreasing tendency, *Hif1a*-cKO significantly reduced fibrosis volumes (**Fig. 1B and C**). As VHL is a master regulator of HIF-α proteolysis, we produced RPE-specific *Vhl-*cKO mice (*Vhl*^f/f^; Best1-Cre^tg/-^) (23, 28) to examine the expected opposing effect at the chronic stage of laser-induced subretinal fibrosis (**Fig. 1D**). As expected, large increases in fibrosis volumes were detected under the *Vhl*-cKO condition.

**Fig. 1.**
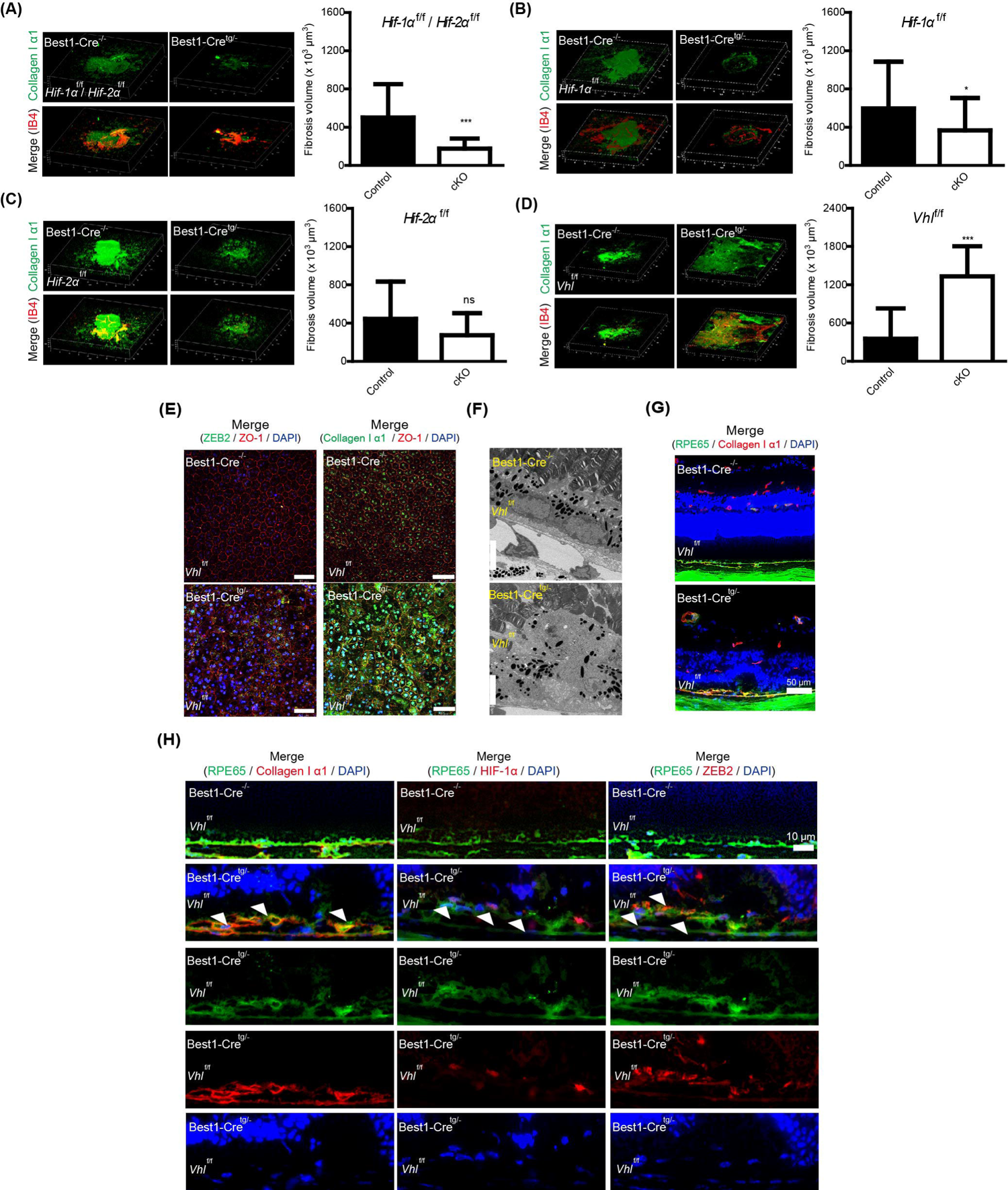
Alterations in subretinal fibrosis by genetic regulation of hypoxia-inducible factors (HIFs). **(A)** Representative images and quantitative data (n = 21–22 per group) demonstrated that significant reductions in fibrosis volumes stained by collagen I α1 were seen in RPE-specific *Hifs* double conditional knock-out (cKO) mice (*Hif-1*α^f/f^/*Hif-2*α^f/f^; Best1-Cre^tg/-^), compared to control mice (Best1-Cre^-/-^). **(B and C)** Representative images and quantitative data (n = 21–54 per group) demonstrated that significant reductions in fibrosis volumes stained by collagen I α1 were seen in RPE-specific *Hif-1*α cKO mice (*Hif-1* ^f/f^; Best1-Cre^tg/-^), compared to control mice (Best1-Cre^-/-^). However, only slight reductions in fibrosis volumes stained by collagen I α1 were seen in RPE-specific *Hif-2*α cKO mice (*Hif-2* ^f/f^; Best1-Cre^tg/-^), compared to control mice (Best1-Cre^-/-^). **(D)** Representative images and quantitative data (n = 19–22 per group) demonstrated that significant increases in fibrosis volumes stained by collagen I α1 were seen in RPE-specific *Vhl* cKO mice (*Vhl*^f/f^; Best1-Cre^tg/-^), compared to control mice (Best1-Cre^-/-^). ns, not significant. *P < 0.05, ***P < 0.001. Two-tailed Student’s *t*-test. Mean ± standard deviation. IB4, Isolectin-IB4. **(E)** Flat-mount images showed that increases in ZEB2 and collagen I α1 expressions were detected in RPE-specific *Vhl* cKO mice (*Vhl*^f/f^; Best1-Cre^tg/-^), compared to control mice (Best1-Cre^-/-^). DAPI, nucleus. Based on the ZO-1 staining image, RPE-barrier dysfunction was further detected. Scale bar: 50 μm. **(F)** Transmission electron microscope (TEM) images showed RPE polarity loss and basal infoldings dysfunction in RPE-specific *Vhl* cKO mice (*Vhl*^f/f^; Best1-Cre^tg/-^), compared to control mice (Best1-Cre^-/-^). Scale bar: 5 μm. **(G and H)** Histological images showed that damages not only in the RPE layer but also in the retinal layer were detected in RPE-specific *Vhl* cKO mice (*Vhl*^f/f^; Best1-Cre^tg/-^), compared to control mice (Best1-Cre^-/-^). Especially, increases in collagen I α1, HIF-1α, and ZEB2 expressions (white arrows in each image) were detected in the RPE layer. Scale bar: 10 μm.

RPE-barrier dysfunction as well as promotion of EMT were present in RPE cells of RPE-specific *Vhl*-cKO mice, even in the absence of laser irradiation, as shown by ZO-1, collagen I α1, and ZEB2 immunostaining (**Fig. 1E**). RPE polarity loss and basal infolding dysfunction were further detected using TEM (**Fig. 1F**). Damage was also detected in the RPE and the retinal layers in RPE-specific *Vhl*-cKO mice (**Fig. 1G**). Increases in collagen I α1, HIF-1α, and ZEB2 expression were also detected in the RPE layer of the *Vhl*-cKO mice (**Fig. 1H**).

### Treatments of HIF inhibitors suppress fibrosis formation in adult mice

We previously found that RPE- or neural retina-specific *Hifs*-cKO and several HIF inhibitors from natural foods (rice bran, vitamin B6, lactoferrin, *G.cambogia*, and hydroxycitric acid) could reduce CNV volumes in a murine model of laser-induced CNV (23, 29, 30). Consistent with this and the hypothesis that HIF-1α could be expressed in laser-induced CNV (**Fig. 2A**), we used general HIF inhibitors (topotecan and doxorubicin) in the same model to examine whether pharmacologic HIF inhibition could be effective in reducing CNV volumes (**Fig. 2B**). Topotecan and doxorubicin treatment significantly reduced CNV volumes. Independent from the CNV volumes, subretinal fibrosis volumes were reduced by treatment with topotecan or doxorubicin (**Fig. 2C and D**). Proteins in the zinc finger E-box binding homeobox (ZEB) family are primarily known as EMT inducers (31, 32), and the expression of the ZEB family proteins was therefore examined under this condition (**Fig. 2E**). Although the expression of *Zeb1* mRNA was not directly upregulated by laser irradiation, that of *Zeb2* mRNA was significantly increased in the mouse RPE-choroid complex. Under the same condition, treatment with topotecan or doxorubicin significantly decreased this mRNA expression in the complex. To support this outcome, a human RPE cell line (ARPE-19) was used in our system (**Fig. 2F**). *COL1A1* mRNA expression was slightly increased under the 1% oxygen hypoxic condition, but topotecan or doxorubicin treatment significantly reduced this upregulation. Furthermore, as with the *in vivo* results, alterations in *ZEB1* or *ZEB2* mRNA expression were detected following topotecan or doxorubicin treatment under the 1% oxygen hypoxic condition in ARPE-19 cells.

**Fig. 2.**
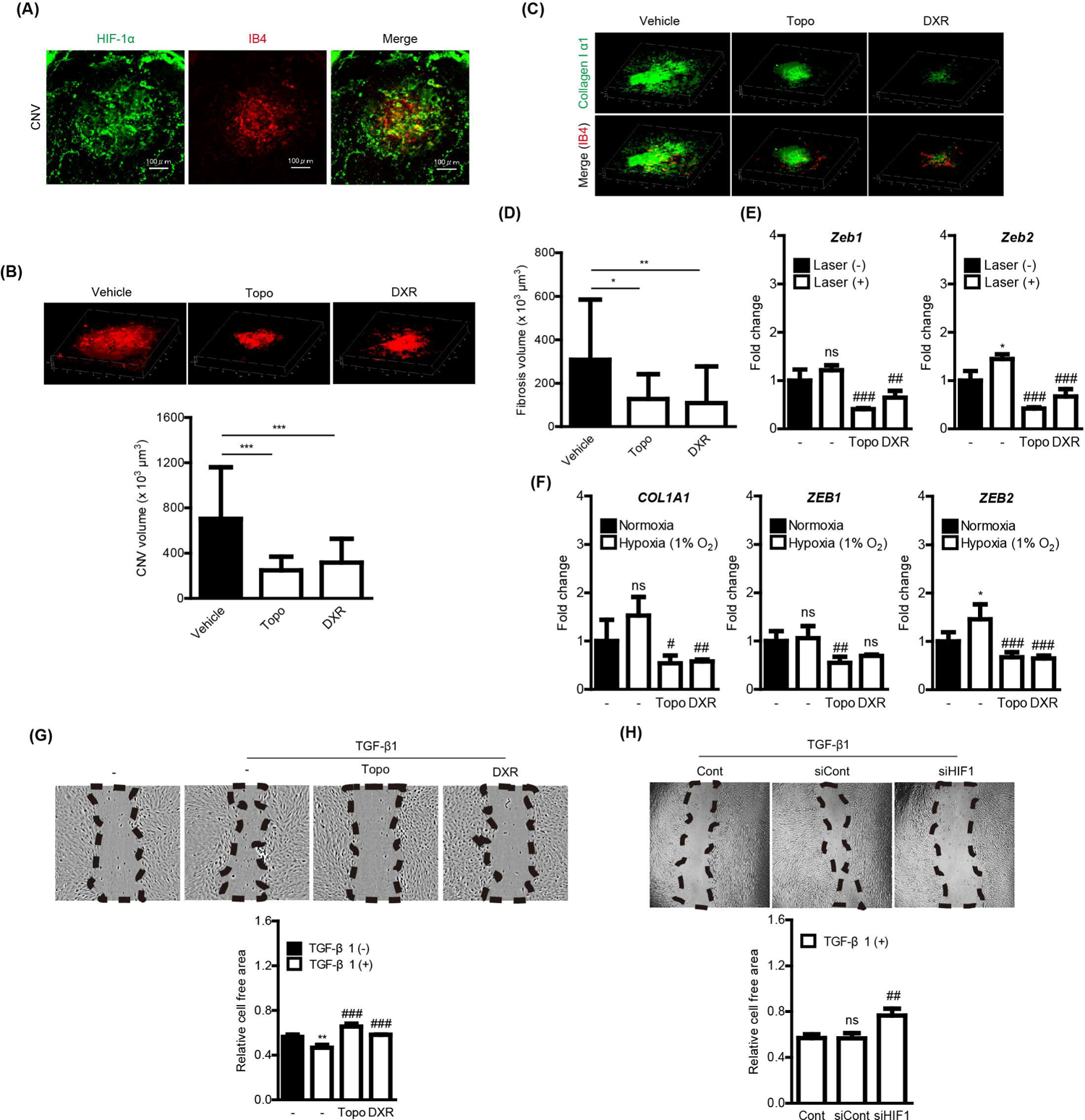
Suppression in subretinal fibrosis by treatments of hypoxia-inducible factor (HIF) inhibitors. **(A)** Immunohistochemistry images showed that HIF-1α was co-labeled with isolectin-IB4 (IB4) in choroidal neovascularization (CNV). **(B)** Representative images and quantitative data (n = 15–26 per group) demonstrated that CNV volume was significantly reduced by administrations of topotecan (Topo) or doxorubicin (DXR). **(C and D)** Representative images and quantitative data (n = 11–24 per group) demonstrated that fibrosis volume stained by collagen I α1 was significantly reduced by administrations of Topo or DXR. *P < 0.05, **P < 0.01, ***P < 0.001. Two-tailed Student’s *t*-test. **(E)** Quantitative data (n = 3 per group) showed that *Zeb1* and *Zeb2* mRNA expressions in the choroid-retinal pigment epithelium (RPE) complex were changed by administrations of Topo or DXR. *P < 0.05 (naïve vs laser irradiation). ##P < 0.01, ###P <0.001 (no drug vs each drug under the laser irradiation condition). One-way ANOVA followed by a Bonferroni post hoc test. **(F)** Quantitative data (n = 3–4 per group) showed that *COL1A1*, *ZEB1*, or *ZEB2* mRNA expression in ARPE-19 cells was changed by treatment of Topo or DXR under the hypoxic condition (1% oxygen). *P < 0.05 (normoxia vs hypoxia). #P < 0.05, ##P < 0.01, ###P <0.001 (no drug vs each drug under the hypoxic condition). One-way ANOVA followed by a Bonferroni post hoc test. **(G)** Representative images and quantitative data (n = 3 per group) demonstrated that reductions in the cell free area by TGF-β1 treatment was significantly suppressed by treatment of Topo or DXR. **P < 0.01 (control vs TGF-β1). ###P <0.001 (no drug vs each drug under the TGF-β1 incubating condition). One-way ANOVA followed by a Bonferroni post hoc test. **(H)** Representative images and quantitative data (n = 3–4 per group) demonstrated that the cell free area was significantly increased by siRNA-mediated genetic HIF inhibition. ns, not significant. ##P <0.01 (siCont; scramble control vs siHIF1; siRNA for *Hif-1*α under the TGF-β1-incubating condition). One-way ANOVA followed by a Bonferroni post hoc test. Mean ± standard deviation. IB4, Isolectin-IB4.

Fibrosis and tissue repair involve a complex process (including RPE proliferation and migration) (33), and a scratch assay was therefore performed to explore the effect of HIF inhibition on the TGF-β1-induced EMT ability of ARPE-19 cells (**Fig. 2G**). While TGF-β1 treatment significantly reduced the cell-free area, topotecan or doxorubicin treatment suppressed this reduction. Furthermore, the *Hif1a* siRNA-transfected group showed a similar outcome in comparison with the control (scramble) siRNA-transfected group under the TGF-β1-incubated condition (**Fig. 2H**).

### Biomolecular features in experimental AMD models are similarly detectable in patients with AMD

In our preclinical study, we identified two possible roles for HIF proteins. HIF-1α (but not HIF-2α) could be an important contributor for the development of MNV and further fibrosis. Among the ZEB family, ZEB2 (but not ZEB1) could be involved in the progression of subretinal fibrosis. In accordance with those findings in mice, the expression of HIF-1α (not HIF-2α), ZEB2, and collagen I α1 was similarly detected in patients with polypoidal choroidal vasculopathy (PCV), one type of wet AMD, or Type 2 MNV (**S. Table 1 and Fig. 3**).

**Fig. 3.**
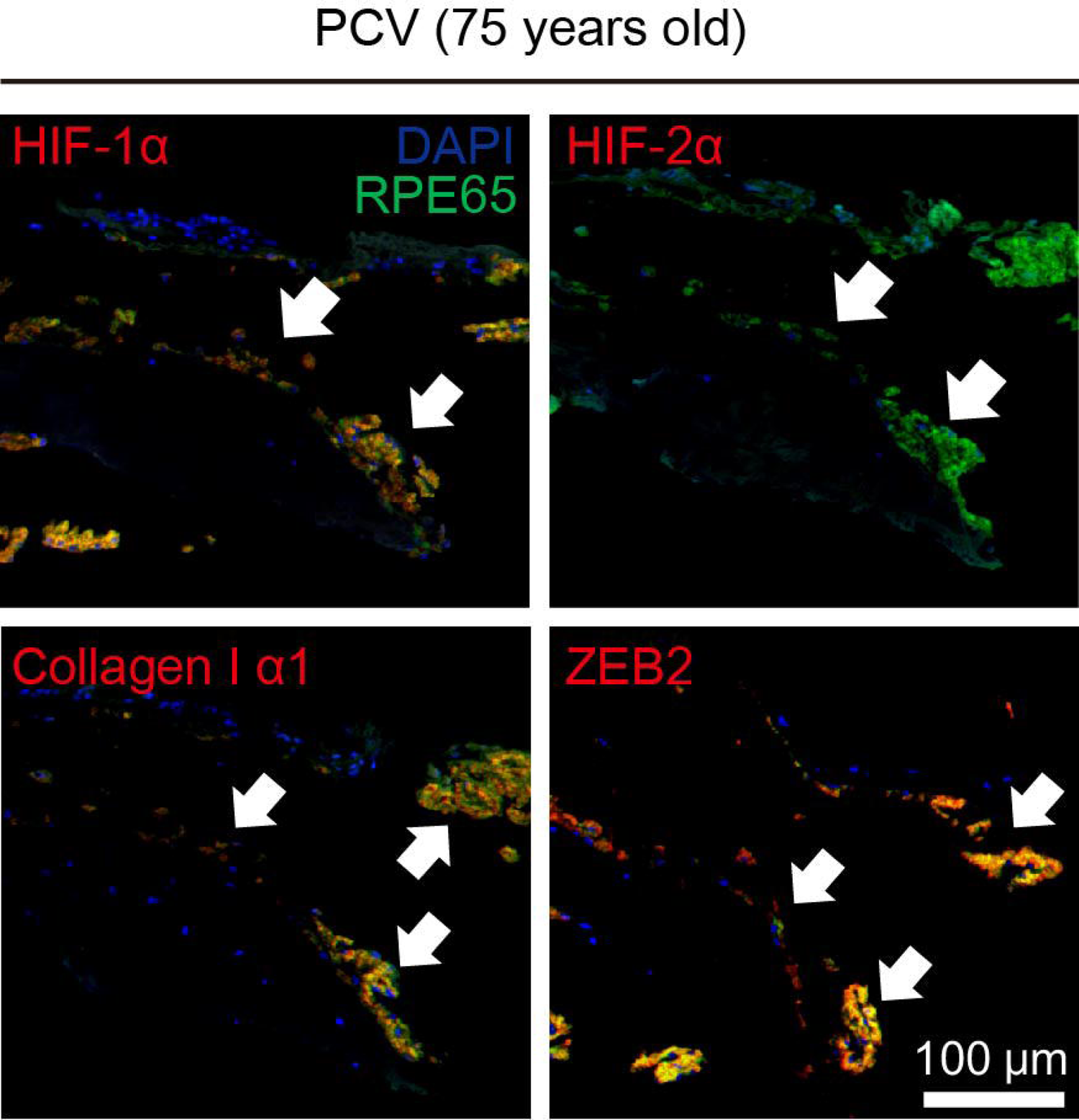
Biomolecular screening of RPE alterations in an AMD patient. Collagen I α1, HIF-1α (not HIF-2α), and ZEB2 expressions (white arrows in each image) were detected in the RPE layer of the AMD patient (the right eye in 75 years old male PCV patient). RPE65 was used to mark the RPE layer. Scale bar: 100 μm.

### Novel HIF inhibitors from marine products suppress fibrosis formation in adult mice

We previously found that extracts from marine products (*D.tabl* and *D.muroadsi*) possessed HIF inhibitory effects under pseudohypoxic and hypoxic conditions (18). We investigated whether those HIF inhibitors could prevent chronic fibrosis formation (**Fig. 4A**). Volumes of fibrosis were considerably reduced by treatment with extracts from these products. As our marine products contained high amounts of histidine and taurine (**S. Table 2**), we investigated whether taurine and histidine could have HIF inhibitory effects in ARPE-19 cells (**Fig. 4B and C**). We found increased HIF-1α expression under the hypoxic condition (1% oxygen) was reduced by treatment with taurine and/or histidine. Furthermore, HIF activities (treatment of DMOG [a competitive inhibitor of HIF-hydroxylated prolyl hydroxylase] or incubation under a 3% oxygen condition) were reduced by treatment with taurine and/or histidine. We then moved to investigate whether retinal neovascularization could be suppressed by taurine or histidine administration (**S. Fig. 1**). Administration of taurine or histidine significantly reduced neovascular tuffs in a murine model of OIR, whereas no changes were evident in vaso-obliteration.

**Fig. 4.**
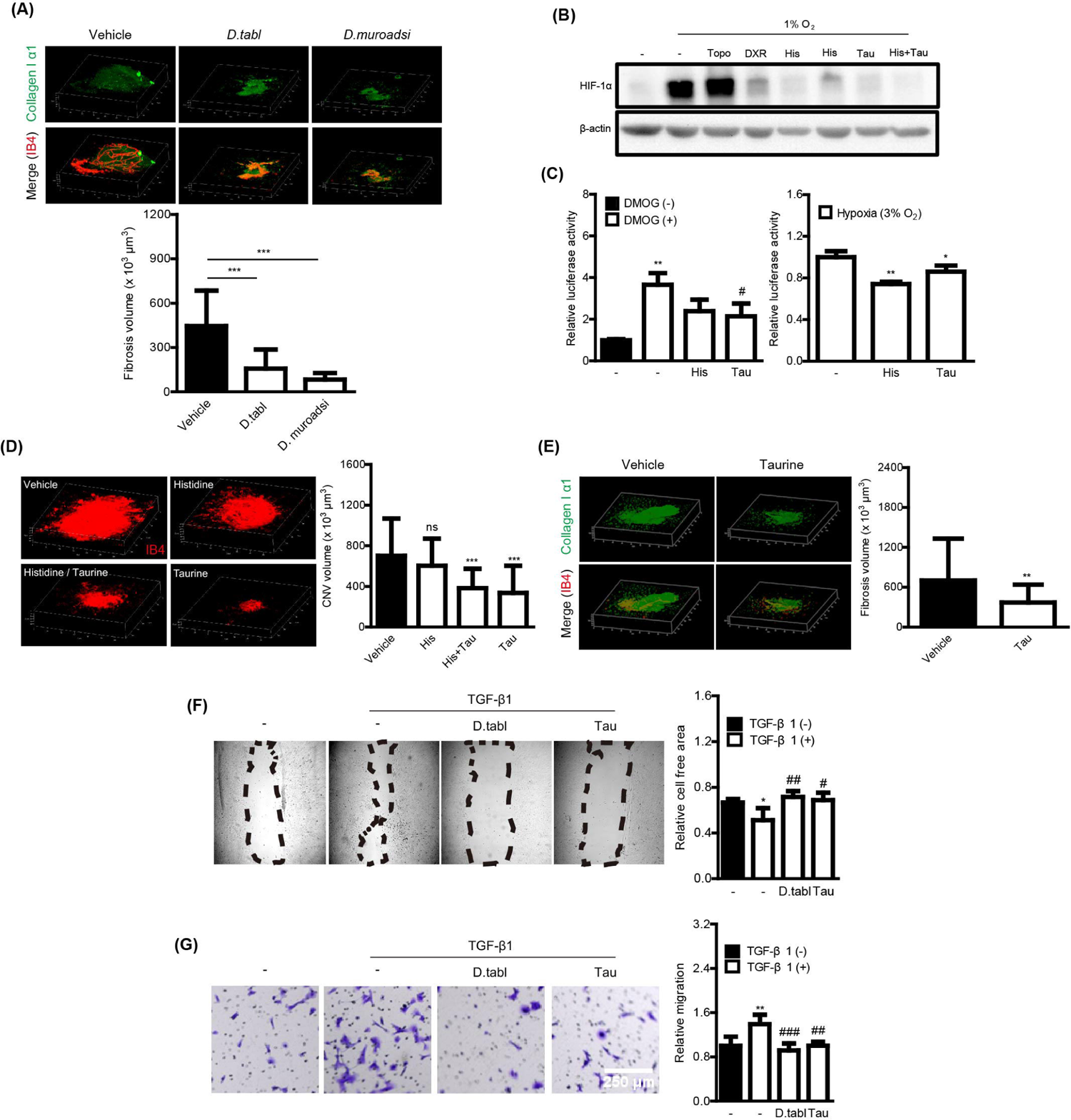
Suppression of subretinal fibrosis by novel HIF inhibitors from marine products. **(A)** Representative images and quantitative data (n = 14–30 per group) demonstrated that fibrosis volumes stained by collagen I α1 were significantly reduced by treatment of extracts from marine products (*D.tabl* and *D.muroadsi*). ***P < 0.001. Two-tailed Student’s *t*-test. IB4, Isolectin-IB4. **(B)** Western blot images showed that increases in HIF-1α expression under the hypoxic condition (1% oxygen) were dramatically reduced by treatment of histidine (His) or taurine (Tau). Topotecan (Topo) and doxorubicin (DXR) were used as generally known HIF inhibitor controls depending on the cell type. **(C)** Quantitative data (n = 3 per group) showed that HIF activation (treatment of DMOG or incubation under the 3% oxygen condition) was significantly reduced by treatment of Tau and/or His. *P <0.05 (no drug vs each drug under the 3% oxygen condition at the bottom), **P < 0.01 (control vs DMOG in the upper part; no drug vs each drug under the 3% oxygen condition at the bottom). #P <0.05 (no drug vs each drug under the DMOG-incubating condition). One-way ANOVA followed by a Bonferroni post hoc test. **(D and E)** Representative images and quantitative data (n = 26–37 per group) demonstrated that choroidal neovascularization (CNV) volumes were significantly reduced by treatment of taurine, while the volumes were not changed by treatment of histidine. Taurine treatment significantly reduced fibrosis volumes at the chronic stage of laser-induced subretinal fibrosis. **P < 0.01, ***P < 0.001. One-way ANOVA followed by a Bonferroni post hoc test; D. Two-tailed Student’s t-test; E. **(F)** Representative images and quantitative data (n = 4–5 per group) demonstrated that reductions in the cell free area by TGF-β1 treatment was significantly suppressed by treatment of extracts from *D.tabl* and taurine (Tau). *P < 0.05 (control vs TGF-β1). ##P <0.01, ###P <0.001 (no drug vs each drug under the TGF-β1-incubating condition). One-way ANOVA followed by a Bonferroni post hoc test. **(G)** Representative images and quantitative data (n = 5 per group) demonstrated that TGF-β1-induced migration was significantly suppressed by treatment of extracts from *D.tabl* and taurine (Tau). Scale bar: 250 μm. **P < 0.05 (control vs TGF-β1). ##P <0.01, ###P <0.001 (no drug vs each drug under the TGF-β1-incubating condition). One-way ANOVA followed by a Bonferroni post hoc test. Mean ± standard deviation

Furthermore, we examined whether taurine and/or histidine could prevent CNV formation in a murine model of laser-induced CNV (**Fig. 4D**). Under this condition, administration of taurine (but not histidine) significantly reduced CNV volumes. Therefore, taurine was further screened for whether its administration could prevent fibrosis formation at the chronic stage, and taurine consequently significantly reduced fibrosis volume (**Fig. 4E**). Furthermore, in the ARPE-19 cell scratch assay, whereas TGF-β1 treatment reduced the cell-free area, treatment with taurine or the extract from *D.tabl* significantly suppressed this reduction; in the migration assay, TGF-β1-induced migration was suppressed by treatment with taurine or the extract from *D.tabl* (**Fig. 4F and G**).

## Discussion

In this study, we demonstrated that pharmacologic and genetic HIF inhibition could reduce the formation of chronic subretinal fibrosis, due to alterations in the EMT inducer gene expression (especially of ZEB2). The in vivo outcomes were further explained using the ARPE-19 in vitro system. Similar patterns of gene expression for *HIF1A* or *ZEB2* were detected in patients with AMD. Importantly, the therapeutic effects of marine products and their major component taurine on the development of fibrosis were found.

Hypoxia is a major pathologic process during ocular ischemia. Under hypoxic conditions as well as abnormal stress conditions, HIF-α is stabilized to translocate into the nucleus, activating the transcription of various HIF-target genes related to angiogenesis, inflammation, and cell death/survival (9). This pathologic process can cause neovascularization in the retina, choroid, or cornea (12). In patients with ischemic retinal diseases, HIF-1α and HIF-2α could promote neovascularization in the retina (34). Among HIFs, HIF-1α has been considered as a main pathologic sensor in the eye (12, 20). A HIF-1 antagonist acriflavine has been shown to suppress ocular neovascularization in murines (35, 36). Digoxin, another HIF-1 inhibitor, inhibited retinal ischemia-induced HIF-1α expression and further suppressed ocular neovascularization (37). A recently developed HIF inhibitor 32-134D (structurally different from acriflavine) was suggested to inhibit the accumulation of HIFs and normalize HIF-target gene expressions in mice and human retinal organoids (38). The accumulation of HIF-1α in response to transient hypoglycemia has also been suggested to worsen ocular conditions in diabetes (39). For the subretinal area, Xie et al. demonstrated that the blockade of the HIF-1α/p53/miRNA-34a axis could reduce laser-induced CNV volumes and mitigate subretinal fibrosis (20), which is similar with our current findings. We also previously found that RPE- or neural retina-specific *Hif1a* cKO could suppress CNV volumes at the acute stage in a murine model of laser-induced CNV (23). Consistent with this, we further found that HIF-1α (rather than HIF-2α) could be an important contributor for subretinal fibrosis by using three different types of HIFs and a type of VHL (a regulator of HIF activities) cKO systems. In our previous study, metabolic defects in the RPE were relatively associated with HIF-2α, rather than HIF-1α (40). Although HIF-2α inhibition did not suppress the fibrosis formation in our current system, additional HIF-2α inhibition under the HIF-1α-inhibiting condition could be considered as double cKO of HIFs more effectively suppressed the fibrosis formation.

As a strong HIF regulator, VHL targets the hydroxylated HIF-α for ubiquitylation and rapid proteasomal degradation under the physiological normoxic condition (41). Dysfunction of VHL could cause HIF-α-mediated upregulation of genes including erythropoietin, VEGF, plate-derived growth factor B, and other glucose uptake- and metabolism-related genes (42). RPE-specific *Vhl*-cKO mice showed pathologic subretinal fibrosis-like features. At the chronic stage after laser irradiation, the large promotion of fibrosis formation was also detected in RPE-specific *Vhl*-cKO mice. Previously, we generated retina-specific cKO mice for *Vhl* and found the arrested transition from fetal to adult circulatory system (28). Deletion of HIF-1α, but not HIF-2α, could rescue vascular defects. Our accumulating data is consistent with disruption of VHL-HIF-1α-further pathologic outcomes.

EMT is a responsive process whereby epithelial cells lose their own properties (such as polarity and cell to cell adhesion), and acquire migratory and invasive features to become mesenchymal cells. In the eye, the EMT has been known to play key roles in the pathogenesis of subretinal fibrosis, the end stage of AMD (43). Previous studies showed that levels of TGF-β1 (one of the most extensively studied EMT inducers (44–46)) were upregulated in mice with subretinal fibrosis in comparison with those on control mice (47, 48). In our current study, TGF-β1-induced EMT was suppressed by pharmacologic and genetic HIF inhibition. TGF has also been detected in tissue sections from patients with AMD, which seems to be related to the development of subretinal fibrosis (49). TGF-β may upregulate the expression of various types of matrix metalloproteinases, downstream contributors of EMT activation (50–53). Although TGF-β-induced immunity and tissue remodeling might be naturally required to some extent, this process could eventually damage retinal tissue structures and induce subretinal fibrosis. Thus, we assume that EMT reduction by modulating HIFs may be a favorable strategy for preventing subretinal fibrosis. ZEB proteins are important transcription factors in the EMT process participating in the numerous life processes, such as embryonic development, fibrosis and tumor progression (54). In colorectal cancer, the promotion of EMT and metastasis has been shown to be involved with HIF-1α via direct ZEB1 modulation (55). Other cancer research demonstrated that the upregulation of ZEB2 by HIF-1α could be associated with the c-Myc pathway, which is important for maintaining the mesenchymal feature of malignant cells (56). Although ZEB1 or ZEB2 has been suggested to contribute to fibrosis development (57–59), in our current study, the expression of ZEB2 was relatively altered under the stress conditions. This expression was modulated by HIF inhibition in RPE cells in vivo and ARPE-19 cells in vitro. Although more studies are needed, the HIF/ZEB2 axis could be involved in the pathologic process of subretinal fibrosis, which needs to be further studied.

Taurine is a non-proteinogenic amino sulfonic acid widely distributed in animal tissues and naturally occurs in foods with protein, such as the meat of fish (60–63). We previously found that marine products (especially, fish) could inhibit HIF activation in ocular cells in vitro and in vivo and further suppress retinal neovascularization in a neonatal murine model of OIR (18). Consistent with this, in our current study, marine products could prevent subretinal fibrosis in a chronic adult murine model of laser-induced subretinal fibrosis. As marine products contain taurine (60), we could reproduce the therapeutic findings using taurine in those two murine neovascularization models. Taurine could inhibit HIF-1α activation under pseudohypoxic and hypoxic conditions in vitro. Based on our migration and scratch assays, taurine could also suppress TGF-β1-induced EMT activation. Until now, although taurine has been reported to be present in the eyes, the main physiological or pathological role is still unclear (62, 64). Reductions in taurine levels were reported during aging in mammals (64, 65). Previous studies demonstrated that supplementation with taurine could improve taurine levels in the eye, increase photoreceptor survival in retinal dystrophy, reduce levels of pro-inflammatory cytokines and chemokines, decrease oxidative stress in the retina, and repair or reduce RPE cell damage (62, 64). Additionally, taurine can improve optic nerve histological alterations, explained with reductions in retinal oxidative stress markers (66). Recently, systemic supplementation with taurine has been suggested to slow various essential markers of aging, such as increases in DNA damage, deficiencies of telomerase activity, dysfunctions of mitochondria, and cellular senescence (67). Age-related metabolic diseases could be consequently associated with loss of taurine in humans. Thus, taurine supplementation could be a good approach to prevent AMD progression.

Researchers have been trying to understand the role of HIFs in various tissues to find a promising cure for ischemic diseases. In ophthalmology, the treatment for ocular neovascular diseases, including diabetic macular edema and AMD, remains repeated injections of anti-VEGF drugs (68). Although this strategy can preserve the visual function, the repeated injections are invasive, cause several complications, and reduce patient adherence to treatment. Failure to respond to anti-VEGF drugs is another important issue (69). HIFs are master regulators of hypoxia-responsive genes including VEGF (70). Accumulating evidence suggests that therapeutics aimed at HIFs might be an alternative, safer, and more effective mode of treatment than only attenuating the amount of VEGF in the treatment of ocular neovascular diseases (71). Therefore, in this study, we sought to identify the pathologic mechanism of ocular ischemia and related promising strategies and suggest the promising HIF inhibition therapy for subretinal fibrosis.

## Methods

### Animals

Adult male mice (C57BL/6J) were purchased from CLEA Japan (Tokyo, Japan). Transgenic mice expressing Cre recombinase under Best1 (Best1-Cre), *Hif1a*-flox, *Hif2a*-flox, and *Vhl*-flox mice were obtained from the Jackson Laboratory (72–75). Transgenic mice with homozygous conditional inactivation of *Hif1* in RPE cells were generated by breeding Best1-Cre transgenic mice with mice bearing the loxP-flanked *Hif1* allele to create first generation *Best1*-Cre, *Hif1a* heterozygous mice (*Hif1a*^f/-^; Best1-Cre^tg/-^). Heterozygous *Best1*-Cre-*Hif1a* mice were subsequently crossed with mice bearing the loxP-flanked *Hif1* allele to obtain homozygous *Best1*-Cre-*Hif1a* mice (*Hif1a*^f/f^; Best1-Cre^tg/-^). Mice with homozygous conditional inactivation of the *Hif1a/Hif2a*, *Hif2a, and Vhl* in RPE cells were generated by the same procedures. The genetic background of all transgenic mice used in this study was C57BL6/J. Cre-negative littermates served as controls in all experiments. After general randomization and 7 days of acclimatization, mice were maintained in a temperature (24 ± 1 °C)-regulated environment (6 mice per cage) under a 12-h light-dark circadian cycle with free access to food and water. During the experimental period, when disease symptoms (e.g., hunched posture, lethargy, undesirable lack of food intake, or unexpected infection) were consecutively observed, a 3× combination of midazolam, medetomidine, and butorphanol tartrate, termed MMB (76), was administered to mice to induce deep anesthesia, and subsequently mice were euthanized. Except for this condition, all mice were used for the current study.

### Patients

Spec imens were surgically extracted from five eyes of five patients who had been diagnosed with neovascular AMD based on fluorescein angiography and clinical findings between 2001 and 2003. These procedures were performed by Nihon University School of Medicine, Japan (77). Surgical excision of subfoveal MNV was performed as previously indicated (78). Surgical specimens were immediately fixed in 10% formalin in phosphate-buffered solution (PBS, pH 7.4) and embedded in paraffin, and serial sectioned slides (4 μm) were prepared.

### Immunohistochemistry

Immunohistochemistry was performed, as previously described (79). O.C.T. compound-embedded blocks containing 4% paraformaldehyde (PFA)-fixed mouse eyes were frozen on dry ice and cut into 12-μm sections at the sagittal plane. After washing with PBS, sectioned samples (human or murine) were incubated overnight with primary antibodies: anti-RPE65 (1:400, Cat# NB100-355, Novus Biologicals, USA), anti-ZEB2 (1:400, Cat# 14026-1-AP, Proteintech, USA), Collagen I α1 (1:400, Cat# NB600-408, Novus Biologicals, USA), anti-HIF-1α (1:400, Cat# NB100-449, Novus Biologicals, USA), anti-HIF-2α (1:400, Cat# NB100-122, Novus Biologicals, USA), anti-ZO-1 (1:400, Cat# 61-7300, Invitrogen, USA). After washing with PBS, secondary antibodies were added to the samples. After washing, DAPI was added to the samples for nuclear staining. After sample mounting, LSM710 microscope (Carl Zeiss, Jena, Germany) was used to visualize immunostaining.

### Cell culture and luciferase assay

The human RPE cell line ARPE-19 was cultured in DMEM/F-12 (Cat #C11330500BT, Gibco, Waltham, MA, USA) media with 10% fetal bovine serum (FBS) and 1% streptomycin penicillin at 37 °C with 5% CO_2_, as previously described (29). Cells were sub-cultured and maintained for the further experiment.

For the luciferase assay, ARPE-19 cells were transfected with a HIF-luciferase reporter gene construct (Cignal Lenti HIF Reporter, Qiagen, Netherlands). The HIF-luciferase construct encodes a firefly luciferase gene under the control of the hypoxia response element; cells were also transfected with a cytomegalovirus-renilla luciferase construct as an internal control. HIF inhibitor screening was performed, as previously indicated (29). To evaluate the inhibitory effect of taurine or histidine, cells were treated with each compound (1 mg/mL) under various pseudohypoxic or hypoxic conditions. After incubation for 24 h at 37 °C with 5% CO_2_, luminescent values were measured using a Dual-Luciferase Reporter Assay System (Promega, Madison, WI, USA).

### Laser-induced CNV and fibrosis

Laser-induced CNV was performed, as previously described (29, 79). Tropicamide and phenylephrine (Santen Pharmaceutical, Osaka, Japan) were applied to induce pupil dilation. A mixture of MMB was injected to induce anesthesia. Subsequently, the eyes of anesthetized mice were gently touched with a contact lens for the preparation of laser irradiation. CNV spots (532 nm argon laser, 100 mw, 100 ms, 75 µm) were made between the retinal vessels at 2-disc diameters from the optic nerve head of mice. Air bubbles were detected as a successful sign of disruption of the Bruch’s membrane by laser irradiation. CNV spots without air bubbles or the spots with unexpected hemorrhage were not included for further analyses. After euthanasia (on week 1 after laser irradiation), micro-scissors were used to obtain flat-mounted choroidal complexes from the eyeballs. Isolectin IB4 solution (IB4 from *Griffonia simplicifolia*, Invitrogen, USA) was used for detecting CNV. A LSM710 microscope (Carl Zeiss, Jena, Germany) was used to locate and visualize CNV. CNV volumes were calculated using the surface and volume tools in Imaris software (Bitplane, Zurich, Switzerland). For fibrosis formation, samples were prepared on week 5 after laser irradiation. Fibrosis was detected using collagen I α1 solution; fibrosis volumes were also calculated using Imaris software. HIF inhibitors topotecan (Topo; 2.5 mg/kg) and doxorubicin (DXR; 1 mg/kg) were intraperitoneally injected to mice once daily for 5 days after laser irradiation for CNV analysis, and Topo (0.625 mg/kg) and DXR (0.25 mg/kg) were injected to mice once daily for 5 days per week after laser irradiation for fibrosis analysis. *Decapterus tabl*, *D.tabl*, (6 g/kg), *Decapterus muroadsi*, *D.muroadsi*, (2.8 g/kg), histidine (Cat# 082-00683, Wako, Japan) (5 g/kg), and taurine (Cat# 203-00111, Wako, Japan) (0.5 g/kg) were orally administered to mice once daily for 5 days after laser irradiation for CNV analysis, *D.tabl* (6 g/kg), *D.muroadsi* (2.8 g/kg), and taurine (0.15 g/kg) were orally administered to mice once daily for 5 days per week after laser irradiation for fibrosis analysis. *D.tabl* and *D.muroadsi* extracts were prepared as previously described (18).

#### Oxygen-induced retinopathy

A murine model of oxygen-induced retinopathy (OIR) was used, as previously developed (80). Postnatal day 8 (P8) mice were placed in an 85% oxygen chamber for 72 h with their mothers. Then, all mice returned to normoxia until P17. Pups received oral administrations of histidine (5000 mg/kg), taurine (500 mg/kg), or ultrapure water as a vehicle from P12 to P16. For the oral administration, a thin tube was inserted into the mouths of pups. At P17, mice were euthanized, and the eyes were excised. Eye samples were fixed in 4% PFA solution, and were flat-mounted. Flat-mounted retinas were stained with Isolectin IB4 solution for 3 days. After encapsulation, neovascular tufts and vaso-obliteration in the retina were observed with a fluorescence microscope (BZ-9000, KEYENCE, Japan). The number of pixels in neovascular tufts and vaso-obliterations was measured in Photoshop (Adobe, San Jose, CA, USA).

#### Quantitative PCR and Western blotting

Quantitative PCR (qPCR) was conducted, as previously described (29, 79). In vitro samples were prepared 12 h after each administration of drug to cells, and in vivo samples were prepared immediately after mice were euthanized depending on the experimental condition. RNA extraction was performed using a RNeasy Plus Mini Kit (Qiagen, Netherlands). cDNA was prepared using a ReverTra Ace qPCR RT Master Mix with gDNA Remover (TOYOBO, Osaka, Japan). Real-time qPCR was performed using a THUNDERBIRD SYBR qPCR Mix (TOYOBO, Osaka, Japan) with the Step One Plus Real-Time PCR system (Applied Biosystems, USA). Information on primers is shown in **S. Table 3**. The fold change between levels of different transcripts was calculated by the ΔΔCT protocol.

Western blotting was conducted as previously described (80, 81). In vitro samples were prepared 48 h after each drug administration to cells. Samples were homogenized on ice in lysis RIPA buffer (Thermo Fischer Scientific, Waltham, USA) containing protease inhibitor cocktail (Roche Diagnostics, Switzerland). Sodium dodecyl sulfate (SDS) loading buffer was added to the samples lysates, which were then heated at 95 °C and separated using 10% SDS-polyacrylamide gel electrophoresis, and transferred to polyvinylidene fluoride membranes. After blocking with nonfat dry milk solution, membranes were incubated with primary antibodies at 4°C overnight: anti-HIF-1α (1:1000, Cat #36169, Cell Signaling Technology, USA) or anti-β-Actin (1:5000, Cat #3700, Cell Signaling Technology). Membranes were washed with TBST three times, and incubated with HRP-conjugated secondary antibodies (1:5000, GE Healthcare, USA). Protein bands were visualized using an ECL kit (Ez WestLumi plus, ATTO, Tokyo, Japan) via chemiluminescence (ImageQuant LAS 4000 mini, GE Healthcare, USA) and quantified using NIH ImageJ software (National Institutes of Health, USA).

#### Transmission electron microscopy

Transmission electron microscopy (TEM) was performed, as previously described (82). Briefly, PBS-perfused eyes were fixed in ice-cold 2.5% glutaraldehyde in 0.1_M_PB (pH_7.4). After washing with 0.1_M_PB, eye samples were post-fixed with 1.0% osmium tetroxide solution (TAAB Laboratories, UK). Samples were then sequentially dehydrated with ethanol (from 50-100%), soaked with acetone and with n-butyl glycidyl ether twice for 30 min with a graded concentration of Epoxy resin with QY-1 for 60 min each, and finally incubated with 100% Epoxy resin (composed of 27.0_g MNA, 51.3_g EPOK-812, 21.9_g DDSA, and 1.1_mL DMP-30; all from Okenshoji Co. Ltd., Tokyo, Japan) with an Automated Tissue Processor (EM-TP, Leica Microsystems, Germany). Samples were polymerized with 100% Epoxy resin at 60 °C in an experimental oven. Resin blocks of each sample were gently shaped, semi-thin sliced to a 2_μm thickness and stained with toluidine blue. Blocks were ultra-thin sectioned to a 80_nm thickness using an ultramicrotome (EM UC7, Leica Microsystems, Germany). Ultra-thin sections were collected onto copper grids and were incubated in uranyl acetate and lead citrate solution. Processed samples were observed using a JEM-1400Plus microscope (JEOL Ltd., Tokyo, Japan).

#### Scratch and migration assays

The scratch assay was performed as described with minor modifications (83–85). After seeding of ARPE-19 cells (at approximately 100% confluence) as a monolayer, wounds were gently made in the center of the well using a pipette tip. After removing cell debris by PBS washing, TGF-β1 (10 ng/mL) was added to cells with each drug condition (1 mg/mL). For siRNA-related experiments, control siRNA (Cat# 4390843, Ambion Inc., USA) and *HIF1A* siRNA (Cat# 4427037, Ambion Inc., USA) were prepared and transfected to ARPE-19 cells according to the manufacturer’s instructions (Lipofectamine RNAiMAX reagent packages, Cat# 13778, Invitrogen, USA). Images of the wound regions were captured 24 h after wound induction. The relative cell-free area was determined by collecting images.

A migration assay was conducted as previously described with minor modifications (86–88). Cells were re-suspended with FBS-free cell medium. The cell suspension was loaded into the upper chamber of each transwell insert (PET membrane 24 well 8.0 µm pore size, Cat# 353097, Falcon, USA), and drug intervention target-containing medium (1 mg/mL) on the lower chamber with TGF-β1 (10 ng/mL) solution. After 9 h of incubation, cells on the upper side of the filter were removed with a cotton swab, and cells that migrated to the lower side were fixed with 4% PFA at room temperature, stained with crystal violet solution (Cat# V5265, Sigma, USA). The relative migration was determined by collecting images.

### Study approval

Whole animal experiments adhered to the Ethics Committee on Animal Research of the Keio University School of Medicine (#16017). The ARVO Statement for the Use of Animals in Ophthalmic and Vision Research, and the international standards of animal care and use, Animal Research: Reporting in vivo Experiments guidelines were also followed.

The Keio University School of Medicine Ethics Committee approved the human study protocol (approval numbers, 202001666). Informed consent for the surgical procedure and for the use of excised tissues was obtained from all patients, in accordance with the tenets of the Declaration of Helsinki.

### Statistics

Statistically significant differences were calculated by using two-tailed Student’s *t*-test or one-way ANOVA followed by a Bonferroni post hoc test depending on the data set written in the each figure legend. P < 0.05 was considered significant.

### Data availability

Data for the current study are available on request from the corresponding author.

## Acknowledgments

We thank K. Kurosaki and A. Kawabata for critical technical supports. The present study was supported by Grants-in-Aid for Scientific Research (KAKENHI, number 15K10881, and 18K09424) from the Ministry of Education, Culture, Sports, Science and Technology (MEXT) to T.K., Funding Support for Development of Research Seeds Based on Marine Biotechnology from Shizuoka Prefecture to T.K., and JST SPRING (number JPMJSP2123) to D.L.

## Contributions

C.S. and D.L. conceived and designed the experiments, performed the experiments, analyzed the data, prepared figures and tables, wrote the first draft of the article, revised the article, and approved the final draft. Y.M. performed the experiments and approved the final draft. H.N. supported the experiments involving human samples. K.Nimura, K.O., S.Y., H.N., H.K., and K.Negishi reviewed and revised drafts of the article and approved the final draft. T.K. supervised the whole study.

## References

1. Mitchell P, Liew G, Gopinath B, and Wong TY. Age-related macular degeneration. Lancet (London, England). 2018;392(10153):1147–59.

2. Grossniklaus HE, and Green WR. Choroidal neovascularization. American journal of ophthalmology. 2004;137(3):496–503.

3. Fleckenstein M, Mitchell P, Freund KB, Sadda S, Holz FG, Brittain C, et al. The Progression of Geographic Atrophy Secondary to Age-Related Macular Degeneration. Ophthalmology. 2018;125(3):369–90.

4. Cheong KX, Cheung CMG, and Teo KYC. Review of Fibrosis in Neovascular Age-Related Macular Degeneration. American journal of ophthalmology. 2023;246:192–222.

5. Ishikawa K, Sreekumar PG, Spee C, Nazari H, Zhu D, Kannan R, et al. αB-Crystallin Regulates Subretinal Fibrosis by Modulation of Epithelial-Mesenchymal Transition. The American journal of pathology. 2016;186(4):859–73.

6. Ishikawa K, Kannan R, and Hinton DR. Molecular mechanisms of subretinal fibrosis in age-related macular degeneration. Experimental eye research. 2016;142:19–25.

7. Zhou M, Geathers JS, Grillo SL, Weber SR, Wang W, Zhao Y, et al. Role of Epithelial-Mesenchymal Transition in Retinal Pigment Epithelium Dysfunction. Frontiers in cell and developmental biology. 2020;8:501.

8. Caceres PS, and Rodriguez-Boulan E. Retinal pigment epithelium polarity in health and blinding diseases. Current opinion in cell biology. 2020;62:37–45.

9. Semenza GL. HIF-1 and mechanisms of hypoxia sensing. Current opinion in cell biology. 2001;13(2):167–71.

10. Walmsley SR, McGovern NN, Whyte MK, and Chilvers ER. The HIF/VHL pathway: from oxygen sensing to innate immunity. American journal of respiratory cell and molecular biology. 2008;38(3):251–5.

11. Ramakrishnan S, Anand V, and Roy S. Vascular endothelial growth factor signaling in hypoxia and inflammation. Journal of neuroimmune pharmacology: the official journal of the Society on NeuroImmune Pharmacology. 2014;9(2):142–60.

12. Lee D, Miwa Y, Kunimi H, Ibuki M, Shoda C, Nakai A, et al. HIF Inhibition Therapy in Ocular Diseases. The Keio journal of medicine. 2022;71(1):1–12.

13. Kurihara T. Roles of Hypoxia Response in Retinal Development and Pathophysiology. The Keio journal of medicine. 2018;67(1):1–9.

14. Kurihara T, Westenskow PD, and Friedlander M. Hypoxia-inducible factor (HIF)/vascular endothelial growth factor (VEGF) signaling in the retina. Advances in experimental medicine and biology. 2014;801:275–81.

15. Zou L, Wang X, and Han X. LncRNA MALAT 1/miR-625-3p/HIF-1α axis regulates the EMT of hypoxia-induced RPE cells by activating NF-κB/snail signaling. Experimental cell research. 2023;429(1):113650.

16. Farjood F, Ahmadpour A, Ostvar S, and Vargis E. Acute mechanical stress in primary porcine RPE cells induces angiogenic factor expression and in vitro angiogenesis. Journal of biological engineering. 2020;14:13.

17. Lai K, Luo C, Zhang X, Ye P, Zhang Y, He J, et al. Regulation of angiogenin expression and epithelial-mesenchymal transition by HIF-1α signaling in hypoxic retinal pigment epithelial cells. Biochimica et biophysica acta. 2016;1862(9):1594–607.

18. Shoda C, Miwa Y, Nimura K, Okamoto K, Yamagami S, Tsubota K, et al. Hypoxia-Inducible Factor Inhibitors Derived from Marine Products Suppress a Murine Model of Neovascular Retinopathy. Nutrients. 2020;12(4).

19. Liu Z, Mao X, Yang Q, Zhang X, Xu J, Ma Q, et al. Suppression of myeloid PFKFB3-driven glycolysis protects mice from choroidal neovascularization. British journal of pharmacology. 2022;179(22):5109–31.

20. Xie L, Wang Y, Li Q, Ji X, Tu Y, Du S, et al. The HIF-1α/p53/miRNA-34a/Klotho axis in retinal pigment epithelial cells promotes subretinal fibrosis and exacerbates choroidal neovascularization. Journal of cellular and molecular medicine. 2021;25(3):1700–11.

21. Zou R, Feng YF, Xu YH, Shen MQ, Zhang X, and Yuan YZ. Yes-associated protein promotes endothelial-to-mesenchymal transition of endothelial cells in choroidal neovascularization fibrosis. International journal of ophthalmology. 2022;15(5):701–10.

22. Cui K, Liu J, Huang L, Qin B, Yang X, Li L, et al. Andrographolide attenuates choroidal neovascularization by inhibiting the HIF-1α/VEGF signaling pathway. Biochemical and biophysical research communications. 2020;530(1):60–6.

23. Ibuki M, Shoda C, Miwa Y, Ishida A, Tsubota K, and Kurihara T. Lactoferrin Has a Therapeutic Effect via HIF Inhibition in a Murine Model of Choroidal Neovascularization. Frontiers in pharmacology. 2020;11:174.

24. Little K, Llorián-Salvador M, Tang M, Du X, O’Shaughnessy Ó, McIlwaine G, et al. A Two-Stage Laser-Induced Mouse Model of Subretinal Fibrosis Secondary to Choroidal Neovascularization. Translational vision science & technology. 2020;9(4):3.

25. Zandi S, Li Y, Jahnke L, Schweri-Olac A, Ishikawa K, Wada I, et al. Animal model of subretinal fibrosis without active choroidal neovascularization. Experimental eye research. 2023;229:109428.

26. Liu Y, Noda K, Murata M, Wu D, Kanda A, and Ishida S. Blockade of Platelet-Derived Growth Factor Signaling Inhibits Choroidal Neovascularization and Subretinal Fibrosis in Mice. Journal of clinical medicine. 2020;9(7).

27. Lai K, Li Y, Li L, Gong Y, Huang C, Zhang Y, et al. Intravitreal injection of triptolide attenuates subretinal fibrosis in laser-induced murine model. Phytomedicine: international journal of phytotherapy and phytopharmacology. 2021;93:153747.

28. Kurihara T, Kubota Y, Ozawa Y, Takubo K, Noda K, Simon MC, et al. von Hippel-Lindau protein regulates transition from the fetal to the adult circulatory system in retina. Development (Cambridge, England). 2010;137(9):1563–71.

29. Ibuki M, Lee D, Shinojima A, Miwa Y, Tsubota K, and Kurihara T. Rice Bran and Vitamin B6 Suppress Pathological Neovascularization in a Murine Model of Age-Related Macular Degeneration as Novel HIF Inhibitors. International journal of molecular sciences. 2020;21(23).

30. Ibuki M, Shoda C, Miwa Y, Ishida A, Tsubota K, and Kurihara T. Therapeutic Effect of Garcinia cambogia Extract and Hydroxycitric Acid Inhibiting Hypoxia-Inducible Factor in a Murine Model of Age-Related Macular Degeneration. International journal of molecular sciences. 2019;20(20).

31. Di Gregorio J, Robuffo I, Spalletta S, Giambuzzi G, De Iuliis V, Toniato E, et al. The Epithelial-to-Mesenchymal Transition as a Possible Therapeutic Target in Fibrotic Disorders. Frontiers in cell and developmental biology. 2020;8:607483.

32. Lamouille S, Xu J, and Derynck R. Molecular mechanisms of epithelial-mesenchymal transition. Nature reviews Molecular cell biology. 2014;15(3):178–96.

33. Szczepan M, Llorián-Salvador M, Chen M, and Xu H. Immune Cells in Subretinal Wound Healing and Fibrosis. Frontiers in cellular neuroscience. 2022;16:916719.

34. Zhang J, Qin Y, Martinez M, Flores-Bellver M, Rodrigues M, Dinabandhu A, et al. HIF-1α and HIF-2α redundantly promote retinal neovascularization in patients with ischemic retinal disease. The Journal of clinical investigation. 2021;131(12).

35. Zeng M, Shen J, Liu Y, Lu LY, Ding K, Fortmann SD, et al. The HIF-1 antagonist acriflavine: visualization in retina and suppression of ocular neovascularization. Journal of molecular medicine (Berlin, Germany). 2017;95(4):417–29.

36. Hackett SF, Fu J, Kim YC, Tsujinaka H, Shen J, Lima ESR, et al. Sustained delivery of acriflavine from the suprachoroidal space provides long term suppression of choroidal neovascularization. Biomaterials. 2020;243:119935.

37. Yoshida T, Zhang H, Iwase T, Shen J, Semenza GL, and Campochiaro PA. Digoxin inhibits retinal ischemia-induced HIF-1alpha expression and ocular neovascularization. FASEB journal: official publication of the Federation of American Societies for Experimental Biology. 2010;24(6):1759–67.

38. Zhang J, Sharma D, Dinabandhu A, Sanchez J, Applewhite B, Jee K, et al. Targeting hypoxia-inducible factors with 32-134D safely and effectively treats diabetic eye disease in mice. The Journal of clinical investigation. 2023;133(13).

39. Guo C, Deshpande M, Niu Y, Kachwala I, Flores-Bellver M, Megarity H, et al. HIF-1α accumulation in response to transient hypoglycemia may worsen diabetic eye disease. Cell reports. 2023;42(1):111976.

40. Kurihara T, Westenskow PD, Gantner ML, Usui Y, Schultz A, Bravo S, et al. Hypoxia-induced metabolic stress in retinal pigment epithelial cells is sufficient to induce photoreceptor degeneration. eLife. 2016;5.

41. Zhang J, and Zhang Q. VHL and Hypoxia Signaling: Beyond HIF in Cancer. Biomedicines. 2018;6(1).

42. Krieg M, Haas R, Brauch H, Acker T, Flamme I, and Plate KH. Up-regulation of hypoxia-inducible factors HIF-1alpha and HIF-2alpha under normoxic conditions in renal carcinoma cells by von Hippel-Lindau tumor suppressor gene loss of function. Oncogene. 2000;19(48):5435–43.

43. Shu DY, Butcher E, and Saint-Geniez M. EMT and EndMT: Emerging Roles in Age-Related Macular Degeneration. International journal of molecular sciences. 2020;21(12).

44. Tretbar S, Krausbeck P, Müller A, Friedrich M, Vaxevanis C, Bukur J, et al. TGF-β inducible epithelial-to-mesenchymal transition in renal cell carcinoma. Oncotarget. 2019;10(15):1507–24.

45. Jennings MT, and Pietenpol JA. The role of transforming growth factor beta in glioma progression. Journal of neuro-oncology. 1998;36(2):123–40.

46. Zhang J, Tian XJ, Zhang H, Teng Y, Li R, Bai F, et al. TGF-β-induced epithelial-to-mesenchymal transition proceeds through stepwise activation of multiple feedback loops. Science signaling. 2014;7(345):ra91.

47. Tosi GM, Orlandini M, and Galvagni F. The Controversial Role of TGF-β in Neovascular Age-Related Macular Degeneration Pathogenesis. International journal of molecular sciences. 2018;19(11).

48. Zhang H, and Liu ZL. Transforming growth factor-β neutralizing antibodies inhibit subretinal fibrosis in a mouse model. International journal of ophthalmology. 2012;5(3):307–11.

49. Wang K, Li H, Sun R, Liu C, Luo Y, Fu S, et al. Emerging roles of transforming growth factor β signaling in wet age-related macular degeneration. Acta biochimica et biophysica Sinica. 2019;51(1):1–8.

50. Gomes LR, Terra LF, Wailemann RA, Labriola L, and Sogayar MC. TGF-β1 modulates the homeostasis between MMPs and MMP inhibitors through p38 MAPK and ERK1/2 in highly invasive breast cancer cells. BMC cancer. 2012;12:26.

51. Fuxe J, and Karlsson MC. TGF-β-induced epithelial-mesenchymal transition: a link between cancer and inflammation. Seminars in cancer biology. 2012;22(5-6):455–61.

52. Hsieh HL, Wang HH, Wu WB, Chu PJ, and Yang CM. Transforming growth factor-β1 induces matrix metalloproteinase-9 and cell migration in astrocytes: roles of ROS-dependent ERK- and JNK-NF-κB pathways. Journal of neuroinflammation. 2010;7:88.

53. Hachana S, and Larrivée B. TGF-β Superfamily Signaling in the Eye: Implications for Ocular Pathologies. Cells. 2022;11(15).

54. Vandewalle C, Van Roy F, and Berx G. The role of the ZEB family of transcription factors in development and disease. Cellular and molecular life sciences: CMLS. 2009;66(5):773–87.

55. Zhang W, Shi X, Peng Y, Wu M, Zhang P, Xie R, et al. HIF-1α Promotes Epithelial-Mesenchymal Transition and Metastasis through Direct Regulation of ZEB1 in Colorectal Cancer. PloS one. 2015;10(6):e0129603.

56. Yoo YG, Christensen J, and Huang LE. HIF-1α confers aggressive malignant traits on human tumor cells independent of its canonical transcriptional function. Cancer research. 2011;71(4):1244–52.

57. Cheng L, Zhou MY, Gu YJ, Chen L, and Wang Y. ZEB1: New advances in fibrosis and cancer. Molecular and cellular biochemistry. 2021;476(4):1643–50.

58. Scott CL, and Omilusik KD. ZEBs: Novel Players in Immune Cell Development and Function. Trends in immunology. 2019;40(5):431–46.

59. de Haan W, Dheedene W, Apelt K, Décombas-Deschamps S, Vinckier S, Verhulst S, et al. Endothelial Zeb2 preserves the hepatic angioarchitecture and protects against liver fibrosis. Cardiovascular research. 2022;118(5):1262–75.

60. Larsen R, Eilertsen K-E, Mæhre H, Jensen I-J, and Elvevoll EO. Bioactive Compounds from Marine Foods. 2013:249–68.

61. Jangra A, Gola P, Singh J, Gond P, Ghosh S, Rachamalla M, et al. Emergence of taurine as a therapeutic agent for neurological disorders. Neural regeneration research. 2024;19(1):62–8.

62. Castelli V, Paladini A, d’Angelo M, Allegretti M, Mantelli F, Brandolini L, et al. Taurine and oxidative stress in retinal health and disease. CNS neuroscience & therapeutics. 2021;27(4):403–12.

63. Jong CJ, Sandal P, and Schaffer SW. The Role of Taurine in Mitochondria Health: More Than Just an Antioxidant. Molecules (Basel, Switzerland). 2021;26(16).

64. Duan H, Song W, Guo J, and Yan W. Taurine: A Source and Application for the Relief of Visual Fatigue. Nutrients. 2023;15(8).

65. Militante J, and Lombardini JB. Age-related retinal degeneration in animal models of aging: possible involvement of taurine deficiency and oxidative stress. Neurochemical research. 2004;29(1):151–60.

66. Nor Arfuzir NN, Agarwal R, Iezhitsa I, Agarwal P, Sidek S, and Ismail NM. Taurine protects against retinal and optic nerve damage induced by endothelin-1 in rats via antioxidant effects. Neural regeneration research. 2018;13(11):2014–21.

67. Singh P, Gollapalli K, Mangiola S, Schranner D, Yusuf MA, Chamoli M, et al. Taurine deficiency as a driver of aging. Science (New York, NY). 2023;380(6649):eabn9257.

68. Xu M, Fan R, Fan X, Shao Y, and Li X. Progress and Challenges of Anti-VEGF Agents and Their Sustained-Release Strategies for Retinal Angiogenesis. Drug design, development and therapy. 2022;16:3241–62.

69. Wallsh JO, and Gallemore RP. Anti-VEGF-Resistant Retinal Diseases: A Review of the Latest Treatment Options. Cells. 2021;10(5).

70. Rattner A, Williams J, and Nathans J. Roles of HIFs and VEGF in angiogenesis in the retina and brain. The Journal of clinical investigation. 2019;129(9):3807–20.

71. Vadlapatla RK, Vadlapudi AD, and Mitra AK. Hypoxia-inducible factor-1 (HIF-1): a potential target for intervention in ocular neovascular diseases. Current drug targets. 2013;14(8):919–35.

72. Iacovelli J, Zhao C, Wolkow N, Veldman P, Gollomp K, Ojha P, et al. Generation of Cre transgenic mice with postnatal RPE-specific ocular expression. Investigative ophthalmology & visual science. 2011;52(3):1378–83.

73. Ryan HE, Lo J, and Johnson RS. HIF-1 alpha is required for solid tumor formation and embryonic vascularization. The EMBO journal. 1998;17(11):3005–15.

74. Gruber M, Hu CJ, Johnson RS, Brown EJ, Keith B, and Simon MC. Acute postnatal ablation of Hif-2alpha results in anemia. Proceedings of the National Academy of Sciences of the United States of America. 2007;104(7):2301–6.

75. Haase VH, Glickman JN, Socolovsky M, and Jaenisch R. Vascular tumors in livers with targeted inactivation of the von Hippel-Lindau tumor suppressor. Proceedings of the National Academy of Sciences of the United States of America. 2001;98(4):1583–8.

76. Miwa Y, Tsubota K, and Kurihara T. Effect of midazolam, medetomidine, and butorphanol tartrate combination anesthetic on electroretinograms of mice. Molecular vision. 2019;25:645–53.

77. Nakashizuka H, Mitsumata M, Okisaka S, Shimada H, Kawamura A, Mori R, et al. Clinicopathologic findings in polypoidal choroidal vasculopathy. Investigative ophthalmology & visual science. 2008;49(11):4729–37.

78. Lambert HM, Capone A, Jr., Aaberg TM, Sternberg P, Jr., Mandell BA, and Lopez PF. Surgical excision of subfoveal neovascular membranes in age-related macular degeneration. American journal of ophthalmology. 1992;113(3):257–62.

79. Lee D, Nakai A, Miwa Y, Negishi K, Tomita Y, and Kurihara T. Pemafibrate prevents choroidal neovascularization in a mouse model of neovascular age-related macular degeneration. PeerJ. 2023;11:e14611.

80. Lee D, Miwa Y, Wu J, Shoda C, Jeong H, Kawagishi H, et al. A Fairy Chemical Suppresses Retinal Angiogenesis as a HIF Inhibitor. Biomolecules. 2020;10(10).

81. Lee D, Tomita Y, Jeong H, Miwa Y, Tsubota K, Negishi K, et al. Pemafibrate Prevents Retinal Dysfunction in a Mouse Model of Unilateral Common Carotid Artery Occlusion. International journal of molecular sciences. 2021;22(17).

82. Kuroha S, Katada Y, Isobe Y, Uchino H, Shishikura K, Nirasawa T, et al. Long chain acyl-CoA synthetase 6 facilitates the local distribution of di-docosahexaenoic acid- and ultra-long-chain-PUFA-containing phospholipids in the retina to support normal visual function in mice. FASEB journal: official publication of the Federation of American Societies for Experimental Biology. 2023;37(9):e23151.

83. Hao XN, Wang WJ, Chen J, Zhou Q, Qu YX, Liu XY, et al. Effects of resveratrol on ARPE-19 cell proliferation and migration via regulating the expression of proliferating cell nuclear antigen, P21, P27 and p38MAPK/MMP-9. International journal of ophthalmology. 2016;9(12):1725–31.

84. Yang J, Li J, Wang Q, Xing Y, Tan Z, and Kang Q. Novel NADPH oxidase inhibitor VAS2870 suppresses TGF-β- dependent epithelial-to-mesenchymal transition in retinal pigment epithelial cells. International journal of molecular medicine. 2018;42(1):123–30.

85. Ma X, Pan L, Jin X, Dai X, Li H, Wen B, et al. Microphthalmia-associated transcription factor acts through PEDF to regulate RPE cell migration. Experimental cell research. 2012;318(3):251–61.

86. Chien HW, Wang K, Chang YY, Hsieh YH, Yu NY, Yang SF, et al. Kaempferol suppresses cell migration through the activation of the ERK signaling pathways in ARPE-19 cells. Environmental toxicology. 2019;34(3):312–8.

87. Li F, Lei C, Gong K, Bai S, and Sun L. Palmitic acid promotes human retinal pigment epithelial cells migration by upregulating miR-222 expression and inhibiting NUMB. Aging. 2023;15(18):9341–57.

88. Li M, Li H, Yang S, Liao X, Zhao C, and Wang F. L-carnitine attenuates TGF- β1-induced EMT in retinal pigment epithelial cells via a PPARγ-dependent mechanism. International journal of molecular medicine. 2021;47(6).

