## Supplemental tables and figures for "Inhibition of hypoxia-inducible factors suppresses subretinal fibrosis"

**Supplementary tables**

**S. Table 1.** Immunohistochemical analyses in AMD patients

| **Diagnosis** | **Age (yr) / Gender / Eye** | **Preoperative visual acuity (LogMAR)** | **Expression (strong +/ weak ± / none -)** |
|---|---|---|---|
| PCV | 72 / Male / Right | 1.0 | HIF-1α (+), HIF-2α (±), ZEB2 (+), Collagen I α1 (+) |
| PCV | 74 / Male / Left | 0.4 | HIF-1α (+), HIF-2α (+), ZEB2 (+), Collagen I α1 (+) |
| PCV | 76 / Male / Left | 0.2 | HIF-1α (+), HIF-2α (+), ZEB2 (+), Collagen I α1 (+) |
| Type 2 MNV | 58 / Male / Left | 0.5 | HIF-1α (+), HIF-2α (±), ZEB2 (+), Collagen I α1 (+) |
| PCV | 75 / Male / Right | 0.6 | HIF-1α (+), HIF-2α (-), ZEB2 (+), Collagen I α1 (+) |

**S. Table 2.** Compositions of histidine and taurine in our marine products

| **Fish** | **Histidine (mg/100 g FW)** | **Taurine (mg/100 g FW)** |
|---|---|---|
| *D.tabl* | 118.0 | 473.6 |
| *D.muroadsi* | 95.2 | 432.1 |

FW, fresh weight.

**S. Table 3.** Primer list

| Name | Direction | Sequence (5' → 3') |
|---|---|---|
| *ZEB1* | Forward | TTACACCTTTGCATACAGAACCC |
| | Reverse | TTTACGATTACACCCAGACTGC |
| *ZEB2* | Forward | GCGATGGTCATGCAGTCAG |
| | Reverse | CAGGTGGCAGGTCATTTTCTT |
| *COL1A1* | Forward | GAGGGCCAAGACGAAGACATC |
| | Reverse | CAGATCACGTCATCGCACAAC |
| *GAPDH* | Forward | TCCCTGAGCTGAACGGGAAG |
| | Reverse | GGAGGAGTGGGTGTCGCTGT |
| *Zeb1* | Forward | GCTGGCAAGACAACGTGAAAG |
| | Reverse | GCCTCAGGATAAATGACGGC |
| *Zeb2* | Forward | CCACGCAGTGAGCATCGAA |
| | Reverse | CAGGTGGCAGGTCATTTTCTT |
| *Gapdh* | Forward | AGGAGCGAGACCCCACTAAC |
| | Reverse | GATGACCCTTTTGGCTCCAC |

**Supplementary figures**

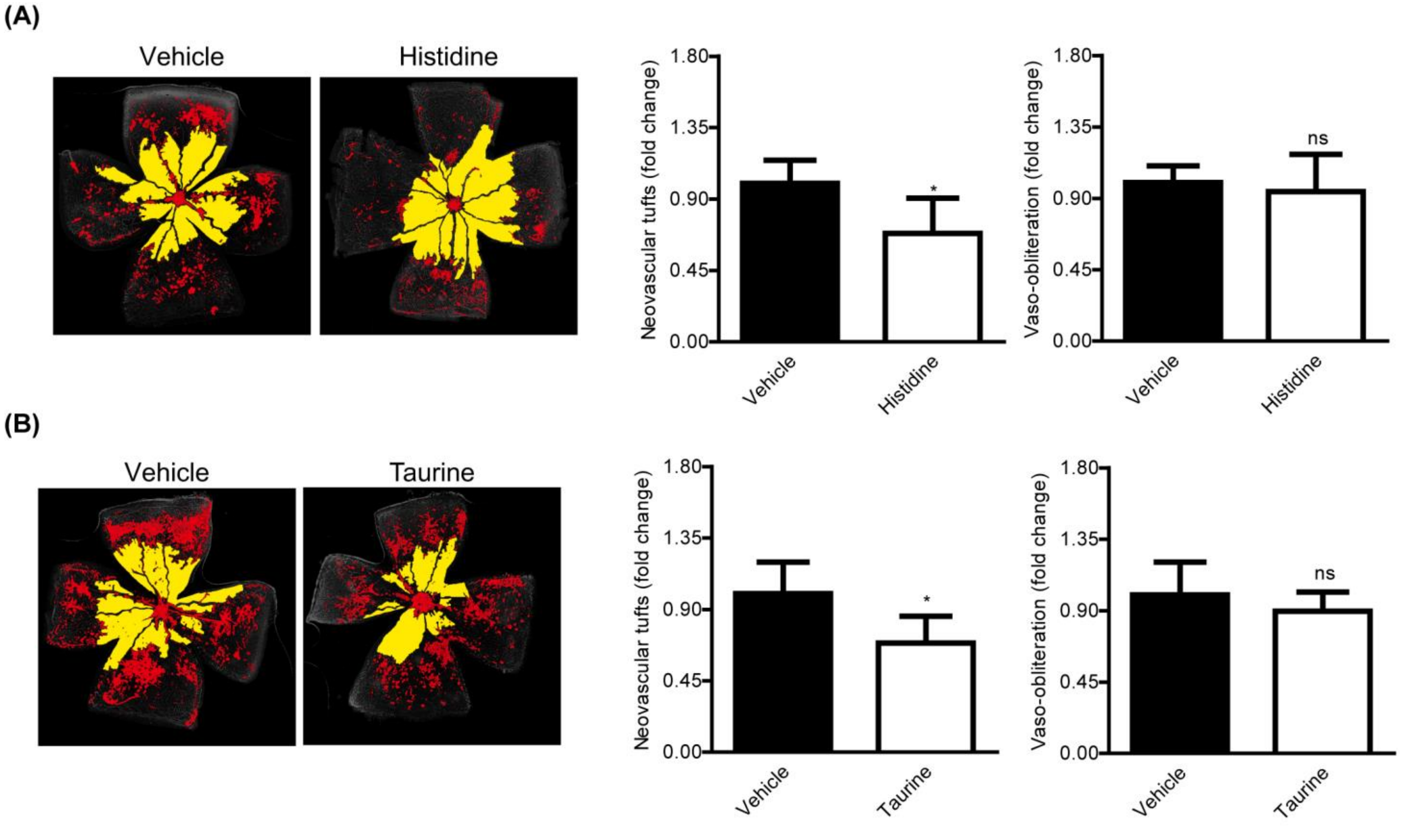


**S. Fig. 1.** Suppression of retinal neovascularization in a murine model of oxygen-induced retinopathy (OIR) by treatment of histidine or taurine. **(A and B)** Images and quantitative data (n = 6 per group) showed that neovascular tufts (red) were reduced by treatment of treatment of histidine (His) or taurine (Tau), while vaso-obliteration (yellow) was not changed. *P <0.05. Two-tailed Student's *t*-test. Mean ± standard deviation.
